# Efficient simulation of neural development using shared memory parallelization

**DOI:** 10.1101/2022.10.17.512465

**Authors:** Erik De Schutter

## Abstract

The Neural Development Simulator, NeuroDevSim, is a Python module that simulates the most important aspects of development: growth, migration and pruning. It uses an agent-based modeling approach inherited from the NeuroMaC software. Each cycle, agents called fronts execute code. In the case of a growing dendritic or axonal front this will be a choice between extension, branching or growth termination. Somatic fronts can migrate to new positions and any front can be retracted to prune parts of neurons.

NeuroDevSim is a multi-core program that uses an innovative shared memory approach to achieve parallel processing without messaging. We demonstrate close to linear strong scaling for medium size models for up to 32 cores and have run large models successfully on 128 cores. Most of the shared memory parallelism is achieved without memory locking. Instead cores have write privileges to private sections of arrays only, while being able to read the entire shared array. Memory conflicts are avoided by a coding rule that allows only active fronts to use methods that need writing access. The exception is collision detection, which is needed to avoid growth of physically overlapping structures. Here a locking mechanism was necessary to control access to grid points that register the location of nearby fronts. A custom approach using a serialized lock broker was able to manage both read and write locking.

NeuroDevSim allows easy modeling of neural development for models ranging from a few complex to thousands of simple neurons or a mixture of both.

## Introduction

Neural development is about more than just growing an organ. During brain development many neurons are born and, in addition, they also establish initial connections. This enables babies to immediately start rewiring their networks correctly during learning. Consequently, studying neural development experimentally is a large domain in neuroscience (Sanes, Reh et al. 2019). Conversely, rather little computational modeling of neural development has been published, despite its clear importance in forming neural networks. The most recent textbook on the subject was published in 2003 (van Ooyen 2003). In other areas of computational neuroscience the development of suitable simulation software has been an important enabler to foster the growth of the subfield. Single neuron modeling became possible with the NEURON (Hines and Carnevale 1997) and GENESIS (Bower and Beeman 1998) simulators, while large scale neural network modeling was greatly facilitated by software packages like NEST (Jordan, Ippen et al. 2018) and Brian (Goodman and Brette 2009). Therefore, it may be useful to do the same for neural development by providing easy to use, computationally performant software that supports modeling multiple aspects of neural development.

Several software packages are available to generate adult dendritic morphologies (Ascoli, Krichmar et al. 2001) or connected networks of neurons with full dendritic morphologies (Koene, Tijms et al. 2009), but these do not claim to simulate the actual developmental sequence. Similarly, we developed a computational framework called NeuroMaC (Neuronal Morphologies and Circuits) that grows multiple neuron morphologies simultaneously, emphasizing the importance of physical interaction between growing neurons (Torben-Nielsen and De Schutter 2014). The only simulator that explicitly models neural development based on intrinsic and extrinsic, contextual factors was CX3D (Zubler and Douglas 2009). It was used to build an impressive simulation of laminated cortex including a few complete neural morphologies (Zubler, Hauri et al. 2013). Unfortunately, it no longer seems to be supported.

This paper presents a new Python software module for Neural Development Simulation, NeuroDevSim, that allows for simulation of most features of neural development. Conceptually it is a successor to the NeuroMaC software, but with greatly improved user interface and computing speed. Software features and several innovative methods to enable pure shared memory parallelization will be described in Materials and Methods. The Results section covers mostly benchmarking of the parallel processing. For those readers familiar with NeuroMaC, comparisons will be made where appropriate.

## Material and Methods

This paper describes NeuroDevSim version 1.0.1 which can be downloaded at https://github.com/CNS-OIST/NeuroDevSim, online documentation is available at https://cns-oist.github.io/NeuroDevSim/index.html. It runs on Python from versions 3.6 onward on MacOS and linux. NeuroDevSim does not work for Windows OS because it requires process forking to implement the shared memory parallelization.

Results were obtained using Python 3.9 either on a MacBook Pro laptop with a M1 Max chip (with 8 performance cores) and 32 Gb memory running macOS Monterey, or on a PC with a 32 core AMD Ryzen Threadripper 3970X and 256 Gb memory running Ubuntu linux version 20. Run times were computed as the difference between start and end times of the *simulation_loop* call. Memory use was tested on the AMD PC.

### Features compared to NeuroMaC

The NeuroMaC software (Torben-Nielsen and De Schutter 2014) focused on modeling the combined influence of internal and external factors on neuron growth, simulated as a sequence of growth *cycles*. The unit of *cycle* is not defined and can correspond to any value in a range of about 1 to 100 hours of real developmental time. Development was embodied in the concept of *fronts* that acted as independent agents simulating growth. A typical sequence was that a new front was made, either as an extension of a neurite or as part of a branching event. On the next cycle this front would be active and execute a neuron specific growth algorithm to decide whether an extension event, a branch event or a growth termination event should occur. The extension event resulted in the creation of one new front, the branching in creation of two or more new fronts and no fronts were created in the case of growth termination. The decision on which event occurred and the position of the new fronts depended on a combination of internal and external factors. Importantly, the software checked for front collisions, new fronts could not overlap with existing ones. At the end of a growth cycle the original front was usually inactivated, it would not grow again. These growth events were stored as a tree structure, where the active front became the parent of one or more child fronts after an extension or branching event.

This functionality is replicated in NeuroDevSim using a different user interface, explained in the next section. Modeling growth is illustrated by replicating the growth of a forest of 100 layer 5 pyramidal neurons (Figure 1A reproduces Figure 4 in <u>(Torben-Nielsen and De Schutter 2014), see also Supp. Movie 1</u>) and the growth of a single alpha motor neuron (Figure1B reproduces Figure 2 in <u>(Torben-Nielsen and De Schutter 2014), see also Supp. Movie 2</u>).

**Figure 1:**
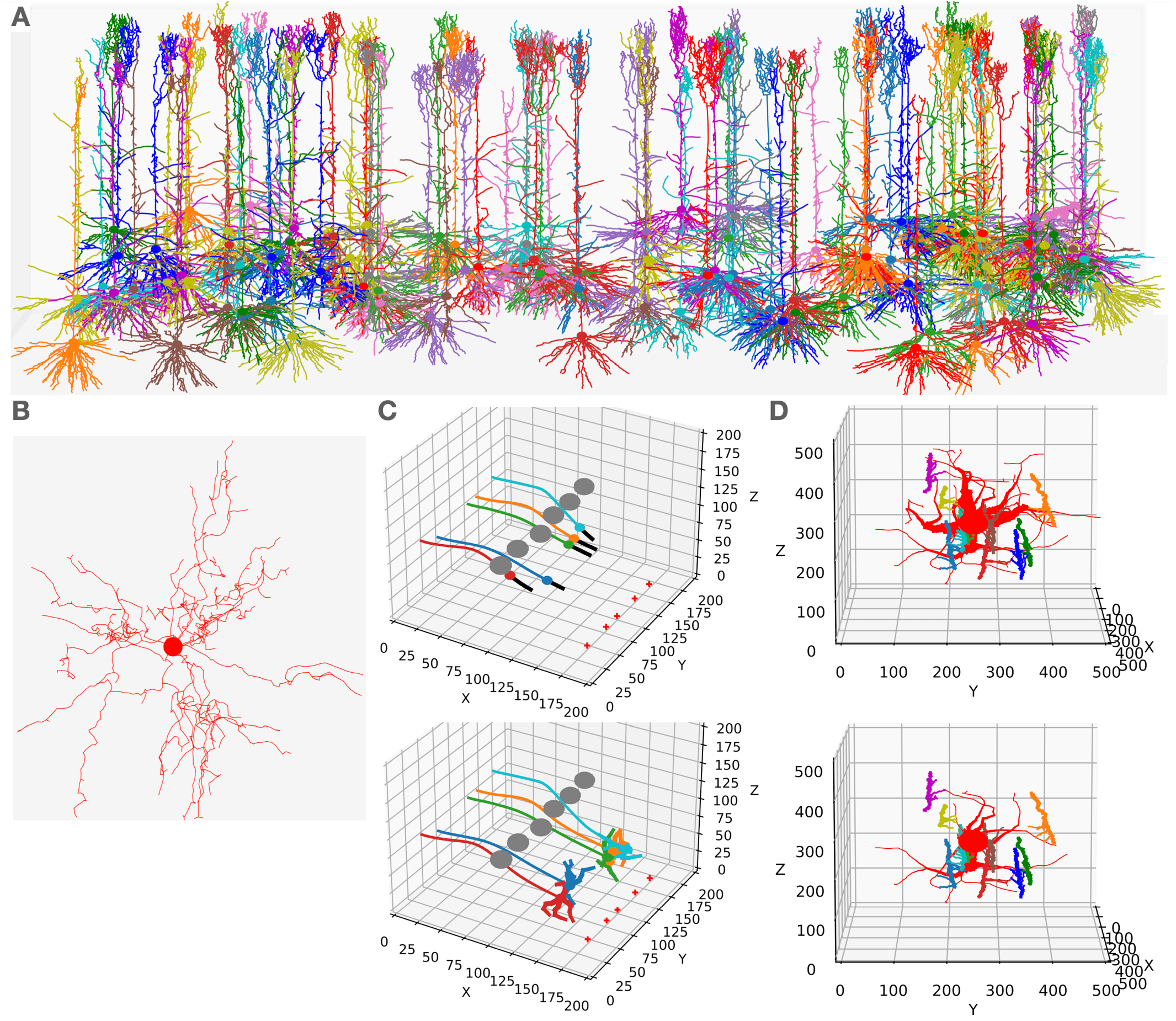
Examples of NeuroDevSim simulations. Panels A-B show results for previously published NeuroMaC simulations and are therefore plotted in the same style: dendrites as wire plots that do not show diameters (with the somata as small circles). Panels C-D are plotted as in the example jupyter notebooks and represent the diameters correctly. All simulations shown are also available as supplementary movies. (**A**) Growth of a forest of 100 layer 5 pyramidal neurons (115 cycles of growth). (**B**) Dendrites of a single alpha motor neuron (71 cycles of growth). (**C**) Migration of 5 somata with leading filipodium and trailing axons. Two images are shown of the simulation: during migration (cycle 20, top) and at end of simulation (cycle 34, bottom). At the bottom right of the simulation volume substrate is placed at 5 locations (red + symbols). The leading filipodia (black color) sense a stochastic representation of the local concentration, based on approximated diffusion of the substrate, and grow towards increasing concentrations. The colored somata migrate along the filipodia and leave behind an axon of the same color. The simulation also contains 6 gray somata that do not grow, but form an obstruction by repelling the growing filipodia. Notice how some of the axons curve around the obstructing gray somata, this shows the path followed by the soma that avoids the obstructions. When the filipodia sense very high substrate concentrations, they retract and migration stops. Note that though the substrate is dispersed along a line, the neurons clearly grew to only two locations and end up forming clusters of two and three neurons respectively. Once migration stops, each soma grows a few dendrites that avoid dendrites of other neurons because no physical overlap is allowed. (**D**) Synapse formation leading to retraction of dendritic branches with few synapses. The red neurons grows 8 dendrites while axons with different colors grow around it from back to front (growth not shown). These axons extend small diameter side branches that make synapses on the red neuron dendrites. The top panel shows the result at cycle 89, before retraction. On cycle 90 five dendrites with less than 5 synapses are retracted instantaneously, the bottom panel shows the result at cycle 91, after retraction. The axon branches making synapses onto the dendrites are more visible in this bottom panel, note that some axon branches now lack a target because no axonal retraction was implemented.

**Figure 2:**
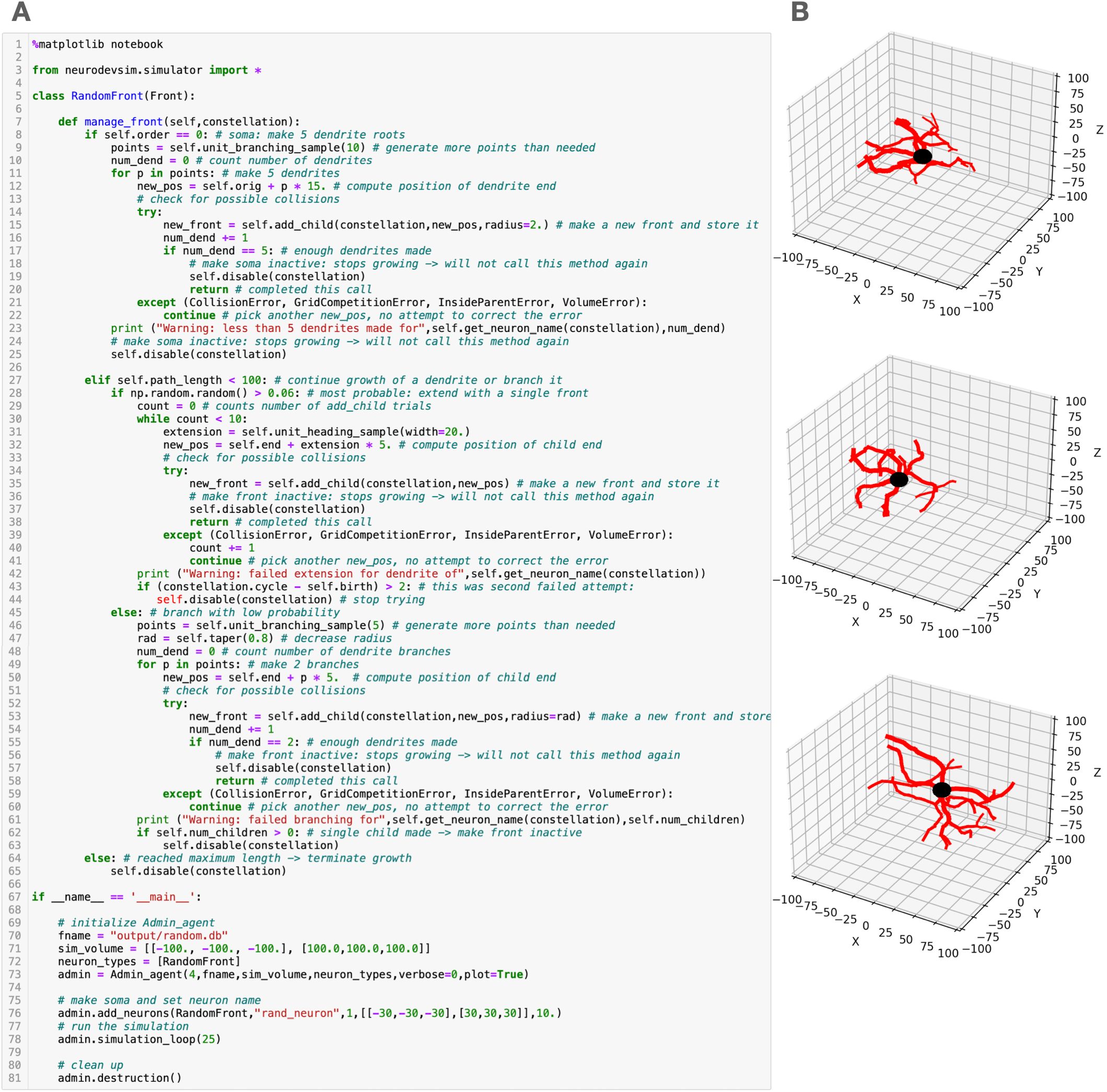
Random growth code example. (**A**) Python simulation script for a very simple NeuroDevSim model, copied from a jupyter notebook. Details of the code are explained in text. (**B**) Three different runs of this script result in very different model results. In these plots the soma is black (default) and dendrite branches are red.

**Figure 3:**
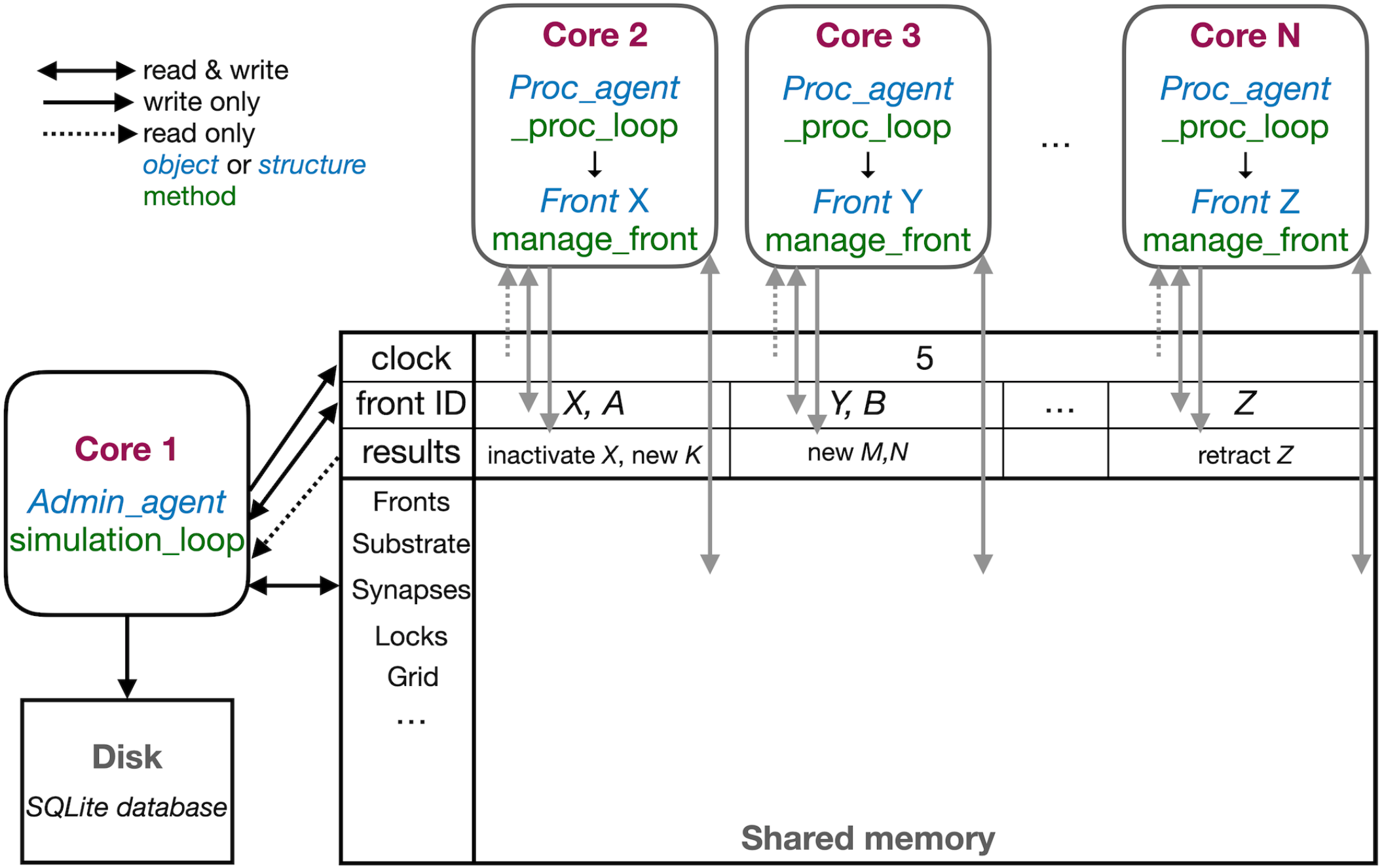
Schematic representation of parallel workflow using shared memory. The default situation, without live plotting, is shown. The simulation is controlled by `Admin_agent` that runs on Core 1 (left). Its interaction with processing cores during execution of `simulation_loop` is shown. `Admin_agent` controls the `clock` that synchronizes all cores and which is available in shared memory. When a new simulation starts, `Admin_agent` puts the front IDs of fronts to be processed in dedicated areas of shared memory for each processing Core 2-N (top). The latter run `_proc_loop` that fetches the front corresponding to the ID, execute its `manage_front` method and puts any new structures in the corresponding arrays in shared memory and their ID plus a symbol in the results section of shared memory. `Admin_agent` then fetches these results to store them in the database (bottom left) and to update its list of active fronts for the next cycle.

**Figure 4:**
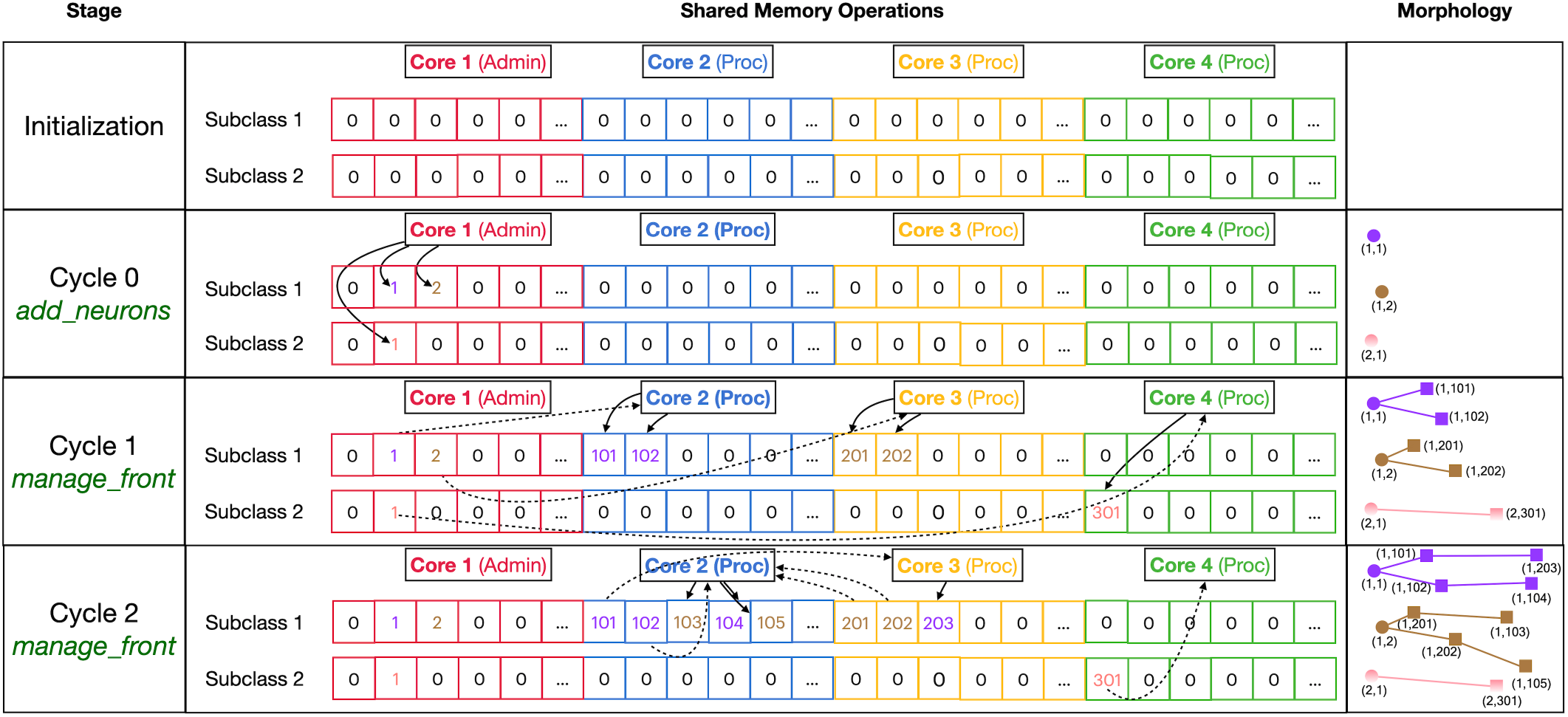
Division of _fronts shared memory array to prevent writing conflicts. The array, shown in the center part, is two-dimensional: each front Subclass has its own row and each Core has its own color coded private section of columns spanning all Subclass rows. Changes in array contents for 4 different stages, indicated in the left part, are shown. The right part of the figure shows schematic 2D representations of the morphology of 3 neurons, two of Subclass 1 and a single of Subclass 2. The fronts are color coded with circles for spherical somata and squares showing `end` of cylindrical fronts. The numbers are front IDs, consisting of (index of subclass array, index in subclass array). In the center sections of the shared `_fronts` array are shown. Zero indicates an empty location, locations that contain a front ID show their index in the corresponding neuron color. Cores can write only to the color coded part allocated to them, full arrows indicate how they write IDs of new fronts at specific indices during each stage. Stippled arrows show how Cores can read front IDs anywhere in the array, so that they can call the corresponding `manage_front` method to make new fronts. See text for more details.

In addition to growth, NeuroDevSim supports simulation of migration and pruning, both important mechanisms in neural development. Migration is limited to somata before they grow dendrites. Two special features can make soma migration more biologically relevant: leading filopodia and trailing axons (Figure 1C). Leading filopodia are growth cone like projections from the soma that can explore the environment to find a proper migration path. NeuroDevSim allows for a single filipodium, with the soma migrating towards the position of the filipodium front that is a child of the soma. The soma can also have an axon, which is automatically extended as the soma migrates.

Pruning is implemented as front retraction, the retracted fronts are removed from the simulation. Entire branches can be retracted in a single cycle (Figure 1D) or slower pruning can be simulated with removal of a single terminal front, possibly followed by more single front retractions during subsequent cycles.

Like in NeuroMaC *substrate* can be used to provide chemical cues to growing fronts. NeuroDevSim has built-in support for approximations of stochastic diffusion (Figure 1C). Where in NeuroMaC synapses were randomly made structures that had only anatomical relevance, in NeuroDevSim synapses are created explicitly during growth (Figure 1D) and can be used to transmit information between neurons at a very long time scale.

### NeuroDevSim user interface

NeuroMaC distributed code and parameters of the Python simulation over several files, including a configuration file. Instead, NeuroDevSim is a Python module that enables description of a complete simulation as a single Python script. This script can be run from a Jupyter notebook, if desired with live graphics, or from the terminal. NeuroDevSim makes use of the Python Errors and Exceptions framework to deal with frequent problems like front collisions and simulation volume borders. An example of a simulation script for a very simple random growth model is shown in Figure 2A and some example results generated in a Jupyter notebook are shown in Figure 2B.

A NeuroDevSim script always contains two components: the definition of one or more `Front` subclasses that describe neuron specific growth rules (lines 5-65 in Fig 2A) and a main program that creates an `Admin_agent`, populates the simulation volume with one or more somata and runs the simulation (lines 67-81 in Figure 2A).

Like in NeuroMaC, the `Admin_agent` object manages the simulation and coordinates the different parallel processes that do the computing, this will be described in more detail in the next section. It also takes care of file output and, if specified, live graphics of the simulation as in this example. The first step in the main program is therefore instantiation of an `Admin_agent` (line 73). Instantiation minimally requires specification of the number of computing processes to be used (4 in the example), the file name of the SQLite (https://www.sqlite.org/) simulation database (`fname`), the simulation volume in cartesian coordinates expressed in μm (`sim_volume`) and a list of all `Front` subclasses to be used (`neuron_types`). Because `Admin_agent` runs on its own core, this model uses 5 cores in total.

Next a number of somata using one of the `Front` subclasses listed in `neuron_types` is created with the `add_neurons` method (line 76). In the example one soma with the neuron name “rand_neuron” is made at a random location within a defined volume and with a radius of 10 μm. In case multiple somata are required, they can either be placed randomly in a volume or on a grid. Somata by default are spheres, other fronts are usually cylinders.

Now the simulation can be run for a fixed number of cycles using the `simulation_loop` method (line 78), which can be called repeatedly so that, for example, additional neurons can be ‘born’ at a later stage. Finally, at the end the `destruction` method should be called to close down all parallel processes and complete the simulation database (line 81).

Several additional methods can be called in the main program, but most of these are beyond the scope of this paper. Importantly, a previous simulation database can be imported immediately after `Admin_agent` instantiation, this can be used to run a series of simulations with a common initial condition.

The algorithms controlling growth etc. are specified in the obligatory `Front` subclass method `manage_front` starting on line 7. Even for this simple `RandomFront`, the `manage_front` definition is fairly long. The `manage_front` method receives as obligatory parameter `constellation` which is a complex data structure, details hidden from the user, that contains all necessary data about the simulation and is passed to many other methods.

As `manage_front` has to deal with all possible `RandomFront` permutations during the entire simulation it is usually structured as a set of nested `if` statements. When the method becomes too long, it can be subdivided by calling specific methods for each condition which improves the readability of the code.

The first `if` statement (line 8) distinguishes between beginning of the simulation when only the soma exists, the growth cycle of dendrites (line 27) and the termination of growth (line 64). The first front created is the soma and this will generate the first call to `manage_front` at cycle 1.

This condition is detected by `if self.order == 0:`, with only somata having a zero order of branching. The soma will grow the roots of 5 dendritic branches, for that purpose 10 random points are generated around the soma with the `unit_branching_sample` method. More points than needed are generated because in a more realistic simulation many cells can be present, resulting in some points already being occupied by other structures. Therefore the dendrite roots are grown by trial and error till either 5 roots are formed or the method runs out of points. This is done by in the loop starting on line 11, which contains a standard sequence to make new fronts.

New fronts will be children of `self`, the front for which `manage_front` is called. For cylindrical new fronts, their first coordinate `orig` will depend on the parent coordinate and only their second coordinate `end` needs to be specified as `new_pos`. For dendrite roots this is a special situation as they are connected to a sphere, which has only one coordinate `orig`. `new_pos` is therefore computed relative to `self.orig`, to which a vector 15 μm long in the random direction *p* is added (line 12). Using this `new_pos` the creation of a new child is tried with the front method `add_child`, note that also a new radius is specified as dendrites should have a smaller radius than the soma. If `add_child` is successful it returns the new front and the number of roots `num_dend` is incremented. If `num_dend` reaches 5, the soma is disabled because it should not grow further and this completes its call of the `manage_front` method (line 20).

Conversely, `add_child` can fail because of a number of errors: the prospective new child may overlap with an existing front causing a `CollissionError`, poorly written code may compute a `new_pos` which is inside the soma causing an `InsideParentError` or which is outside of the simulation volume causing a `VolumeError`, or a `GridCompetitionError` explained in a later section may occur. All of these errors will just cause the code to try the next random point in `points` (lines 39-41). Alternatively, more complex responses to some of these errors can be programmed, for example, the `solve_collision` method may be called to try to overcome a `CollissionError` automatically. This use of the Python Errors and Exceptions framework results in a very flexible and readable way to deal with the problems that naturally arise when simulating growth in a crowded environment using parallel programming.

As mentioned before, the first part of the `manage_front` code is called once when only the soma is present. The next part, for the condition `elif self.path_length < 100:` (line 27) will be called many times and codes most of the model growth. It first makes another decision using an `if` statement: whether to branch or just extend (line 28). This is done by drawing a random number using the `numpy random` library with a 94% probability of extension. For extension a single front will be created. First a pseudo-random direction is drawn that falls within 20 degree cone around the current heading with the `unit_heading_sample` method (line 31). This is then added to the `self.end` coordinate of the parent cylinder with a length of 5 μm. Once again the `add_child` method is tried and errors are caught. If `add_child` is successful the parent is disabled and `manage_front` returns (line 38), otherwise the *while* loop (line 30) is continued and a new pseudo-random direction is drawn. This is repeated till `add_child` is successful, which for this simple model will always work within a few trials.

With 6% probability a branching event will take place. This is done similarly as the code for making dendritic roots at the soma, the difference is that now only two branches need to be made so less random points are requested and a new smaller radius is computed relative to the radius of the parent with the *taper* method (lines 45-63).

Finally, when `self.path_length`, the accumulated distance between the `end` coordinate of `self` and the soma, is longer or equal to 100 μm growth terminates: the front is disabled without calling `add_child` (lines 64-65).

This simple code shows the simplicity of programming robust growth in NeuroDevSim. The full description of the simple random growth model is 64 lines of code. For comparison, the code for the realistic single alpha motor neuron model (Figure 1B) is not much longer at 81 lines with the main differences being branch order dependent control over branching probability and different lengths of individual branches. In contrast, the code for the layer 5 pyramidal neuron forest (Figure 1A) is more complex, with many additional methods, and comprises 372 lines. This is because four different types of dendrites are grown: basal, apical, oblique and tuft dendrites and growth of the latter ones is cortical layer dependent. In addition, repulsion by nearby dendrites from the same neuron is simulated.

The example in Figure 2 uses the `add_child` method only, other NeuroDevSim features are accessible with the front `migrate_soma` (used in Figure 1C), `retract` and `retract_branch` (used in Figure 1D) methods. All of these methods and options to have environmental control of growth are demonstrated in numerous example notebooks that come with the NeuroDevSim distribution.

### Collision detection

A major computing cost is the detection of overlap between existing fronts and new fronts that `add_child` tries to create, such a structural overlap is called a collision. NeuroDevSim uses the same algorithm as NeuroMaC to detect collisions, but in a more scalable implementation. Cylindrical fronts are represented by the line segments that form their central long axes and the distance between the closest points on each line segment is computed using the dist3D_Segment_to_Segment algorithm (http://geomalgorithms.com/). If this distance is smaller than the sum of the radii of the two fronts a collision is detected and `add_child` returns a `CollissionError`.

Obviously, the collision detection should occur only with existing fronts that are spatially close to the proposed new one, as computing distances to all existing fronts rapidly becomes very time consuming. In NeuroMaC, spatial subvolumes partitioned fronts according to their point of origin and collision detection was only performed with fronts in the same subvolume. But as subvolumes could be quite large this often included testing distant fronts.

NeuroDevSim uses a grid to test collisions with close by fronts only. The 3D grid spans the entire simulation volume and grid points are separated by a predefined distance, default 20 μm along all axes. Each front is allocated to at least one grid point, if it is large or positioned about equal distance between multiple grid points it will be allocated to additional grid points. Similarly, the proposed new front is matched to one or more grid points, here called G. Collision detection then involves testing all fronts allocated to G and to their immediate neighboring grid points, with 26 neighbors for each grid point. Standard approach is to raise a `CollissionError` when a first collision is found, with the `CollissionError` returning this first colliding front. It is possible to force a complete search and obtain a list of all colliding fronts, but this slows down the simulator.

### NeuroDevSim code overview

NeuroDevSim uses a very different way to parallelize the simulation than NeuroMaC did. NeuroMaC subdivided the simulation volume into subvolumes, with each spatial subvolume being managed by a different computing core. In addition, a dedicated core ran `Admin_agent` and all front updates had to be communicated to the `Admin_agent` core. Communication between cores was messaging based and implemented with the ZeroMQ (https://zeromq.org) library. This approach to parallelism was simple but created several problems. First, if the number of fronts diverged between subvolumes the simulation became unbalanced with some cores finishing much faster than others. While unbalancing sometimes could be reduced by choosing the right subvolume division scheme, for example not subdividing the vertical axis in the pyramidal neuron forest example (Figure 1A), it led to a limited scaling behavior for many models. Moreover, dealing with fronts that originate in one subvolume but end in another one required communication between cores, that also had to implement the eventuality that such a new front collided with existing ones and had to be discarded. Robust simulation of front migration and retraction across subvolumes was even more challenging.

Conversely, NeuroDevSim does not use spatial subdivision, but instead dynamically allocates fronts to different cores for computation. This allows for a close to perfect balancing between the different computing cores. Moreover, instead of messaging, it uses a shared memory approach that gives each core access to all data and that is also used to schedule jobs for computing cores. The use of shared memory is shown schematically in Figure 3. In this section the general flow of simulating a cycle will be described. The next section will consider parallelization using only shared memory in more detail.

The different cores running `Admin_agent` and, for processing, `Proc_agent` are not synchronized except at the beginning of a cycle. `Admin_agent` controls the cycle, the value of which is stored in shared memory as `clock` so that it can be read by all cores. In contrast with Python practice, everything is numbered from 1, so `Admin_agent` runs on Core 1, because index 0 indicates an unused or empty index in NeuroDevSim. In most arrays index 0 is therefore never used.

Each of the processing cores, numbered Core 2 to N in Figure 3, continuously runs a simple loop `_proc_loop`. It reads the current value of `clock` in shared memory and when the cycle is incremented it starts fetching front IDs from a private section in shared memory. A front ID is an object encoding indexes into the Fronts array in shared memory. The front is accessed, the ID is cleared to signal `Admin_agent` that front processing started and the front’s `manage_front` method is called. For example, in Figure 3 Core 2 receives an instruction to start processing front *X* and it will call the `manage_front` method of *X*. If new fronts or other structures like substrate or synapses are created during `manage_front` they are immediately stored in shared memory, in the Core 2 example a front *K* was created. The outcome of `manage_front` is encoded in the private results section of shared memory as a series of front IDs plus symbols specifying what happened to each front. Note that run times can be very different for each front, this depends on front specific rules and on whether collisions happen or not. In the example shown in Figure 2, the soma front will take more computing time to create 5 dendritic roots than dendrite fronts that make only 1 to 2 children. Also, checking for collisions will be much faster when few fronts are present compared to more crowded areas of the simulation volume. `_proc_loop` keeps fetching front IDs till it receives a unique one that signals end of the cycle. It then performs minimal maintenance and goes into an intermittent sleeping mode till the next cycle starts.

`Admin_agent` controls the cycle, incrementing it from 0 to 1 at the start of the simulation. As mentioned, this signals processing cores to start computing. One task of `Admin_agent` is to supply processing cores with front IDs as fast as possible, ensuring that each core is kept busy. As long as many active fronts need to be computed, it will provide each computing core with 2 front IDs to make sure that the core can run continuously, as shown for Cores 2 and 3 in Figure 3. However, at the end of the cycle, when only a few fronts remain to compute, only a single front ID is provided (Core N in Figure 3). Once all active fronts have been scheduled, the unique end of cycle ID is sent to the core. So at the end of the cycle some processing cores may be idle while others are computing the last fronts for this cycle. The order in which fronts are computed is managed dynamically to avoid grid competition as explained in the next section, in general growing fronts are computed first, followed by migrating ones and then other active fronts. Importantly, the scheduling of front processing to specific computing cores tends to vary for every run of the model because how soon a computing core finishes depends on its working condition, which is beyond the control of NeuroDevSim. Optimal front scheduling requires the user to actively control the growth/migration/active status of fronts. In many cases this just requires calling the `disable` method at proper times, as shown in Figure 2, but more complex code may be required.

Besides scheduling fronts to be computed by processing cores, `Admin_agent` also performs file storage. It uses the information provided by the processing core in their results shared memory to store information about new fronts and other new structures, about soma migration to new positions and about front retractions in the simulation database. Finally, if `simulation_loop` is not finished, it updates its lists of growing, migrating and active fronts to prepare for the next cycle and increments the cycle.

In case of interactive plotting in Jupyter notebooks a second admin core is used and the functionality of `Admin_agent` is split over two cores to improve processing speed. The main admin core still sets the cycle and processes the results returned by processing cores both for file storage and plotting. It also starts the second admin core which only performs front scheduling. A complication is that the main admin core needs to send the second core updated lists of growing, migrating and active fronts at the start of each cycle, which is done through shared memory.

### Parallel computing based uniquely on shared memory

NeuroDevSim uses the `multiprocessing sharedctypes` Python libraries to provide parallelism. All shared data is stored in `RawArrays`, plus `clock` as a `RawValue`. These shared arrays are created early during `Admin_agent` initialization, before the different processing cores are forked. All shared objects, like fronts, substrate and synapses, are defined as `ctypes Structures`. As a consequence their attributes are typed and, because `Structures` are fixed size, instance specific attributes are not supported. However, in the specification of a `Front` subclass the user can define additional attributes that will be present in every instance of the subclass (not shown in this paper).

Because all entries in arrays are fixed size, custom code is needed to deal with Python lists. For short lists, like the list containing the indices of each child of a front, linked lists are used (https://en.wikipedia.org/wiki/Linked_list). For longer lists, like those used by the 3D grid, linked blocks are used. These are accessed by an index to a block of predefined size in the array. Empty spots in the array block can be detected by their zero values and they are typically filled from begin to end. The last entry of each block is an index to a next block in the same array if needed.

Passing simulation data through shared memory is very efficient and simple as long as it is read only. The challenge is to write data in such a way that different cores do not write to the same memory location at the same time. Python provides a `Lock` mechanism, but this does not scale well for the large shared arrays used by NeuroDevSim. Instead specific solutions were developed that avoid locking as much as possible.

The main approach to avoid locking is to subdivide all main data arrays in shared memory into sections so that processing cores write data to their private section only. Figure 4 shows the shared data structure containing Fronts for a fictional example (no code shown) in detail, similar approaches are used to store substrate, synapses, etc. The `_fronts` array is two-dimensional, it is first subdivided by front subclass with in Figure 4 a row for each of 2 subclasses (defined in `neuron_types`). Because all structures are fixed size, and Front subclasses may have different sizes if additional attributes are defined, each subclass is stored in a separate array. These arrays are then subdivided into `num_procs` + 1 equal private sections (colored differently in Figure 4). All cores can read anywhere from these arrays, but only write to their private section which contains 100 entries in this example. Each core maintains counters for the next free index in their private section of each of the subclass specific arrays. Whenever a core makes a new front it checks whether the counter is larger than its maximum index and generates an `OverflowError` if that is the case. Otherwise, it stores the front at counter location and increments the counter.

Figure 4 shows the sequence of the creation of three neurons, two belonging to subclass 1 and a single one for subclass 2 during initiation and the first two cycles of simulation, in a simple case with just 3 processing cores (Cores 2-4). The right column shows a schematic representation of their morphologies in 2D, with circles representing spherical somata and squares representing the `end` coordinate of cylindrical fronts. An `add_neuron` call during initialization (Cycle 0) results in the creation of 3 somata, which will all be stored in the private section of Core 1 which runs `Admin_agent`, each soma in the correct subclass array. Their front IDs are (index of subclass array, index in subclass array) and are shown below the corresponding circles in the Morphology column: (1,1), (1,2) and (2,1). Note that the private section of Core 1 will contain somata only.

During Cycle 1 each of these somata will be processed by different cores: soma (1,1) of the purple neuron is processed by Core 2, soma (1,2) of the brown neuron by Core 3 and soma (2,1) of the pink neuron by Core 4. For each the corresponding core calls their `manage_front` method, which in turn calls `add_child` twice for the purple and brown neurons (subclass 1) and once for the pink neuron (subclass 2). Because the purple neuron was allocated to Core 2, its children are stored in its private section and receive IDs (1,101) and (1,102). Similarly for the brown neuron on Core 3 with IDs (1,201) and (1,202) and the pink one on Core 4 with ID (2,301). Next, during Cycle 2 the children of created during the previous cycle will be processed by different cores. Now the order of scheduling fronts to cores becomes relevant, Figure 4 shows a possible sequence but this could be different in another run of the same model as it depends on in which order new fronts were processed by `Admin_agent` during the previous cycle. The first front to get scheduled is brown front (1,201) for Core 2, this makes a single child (1,103), next is purple front (1,101) on Core 3 making a single child (1,203), followed by pink front (2,301) on Core 4 which does not make a child. The processing of (1,101) on Core 3 and (2,301) on Core 4 takes more time so that they are not available for the remaining two fronts which are, therefore, both processed on Core 2: purple front (1,102) makes child (1,104) followed by brown front (1,202) making its child (1,105).

Note that the entry of the parent, child and soma indices in new fronts, never causes writing conflicts. The parent and soma indices are set in the newly created fronts that are stored in the core’s private section. Child indices are set for the parent front that may be located in another private section (e.g. front (1,101) making child (1,203)), but the parent is the front calling `manage_front` as `self` and during this call it is guaranteed that no other cores will try to change its attributes. Users are instructed not to change front attributes other than those of `self`, except under well defined exceptional circumstances. To enforce this rule, methods like `add_child`, `migrate_soma`, `retract`, etc. raise a `NotSelfError` when not called by `self`.

The approach described combines simplicity with robustness for safely storing new fronts, but results in wasteful use of memory, e.g. in the example of Figure 4 most of the subclass 2 array is not used. Because the private sections of the arrays must be sufficiently large, the overall size of `_Fronts` is large and it usually contains many empty locations. However as this needs to be allocated only once in shared memory (and not separately for every core) the overall memory use is modest for modern architectures (see Results).

However, this approach does not work when fronts are retracted. For simplicity they are not deleted from the `_Fronts` array, so that computing cores do not need to worry about filling empty spaces. Instead they are flagged as retracted and removed from their parent’s list of children. The latter could cause memory writing conflicts when a complete branch is retracted, because some to be retracted fronts at the tips of the branch may simultaneously be calling `manage_front` on another core. Therefore, and as it is assumed that front retractions are rare events, the memory updates needed for front retraction are done in a serial fashion by `Admin_agent` at the end of each cycle.

A major challenge caused by the use of shared memory is the updating of the 3D grid used to detect collisions (Figure 5, top). The grid is mapped onto a one-dimensional integer `_grid` array so that the x,y,z position of grid points uniquely determines an index called *gid*, larger than zero, in this array (Figure 5, mid). Entries in `_grid` are zero for points with no fronts or an index to linked blocks of default size 10 in a `_grid_extra` array containing front IDs (Figure 5, bottom). New blocks in `_grid_extra` are allocated to core specific private sections, but once allocated all cores can write to them. At present, new grid entries are added by traversing the `_grid_extra` block till an empty spot is found. When a grid entry needs to be removed after front retraction, it is replaced by the last entry in `_grid_extra`. Both `_grid` and `_grid_extra` are read and updated frequently by all computing cores, so preventing simultaneous access by different cores has to be controlled carefully.

**Figure 5:**
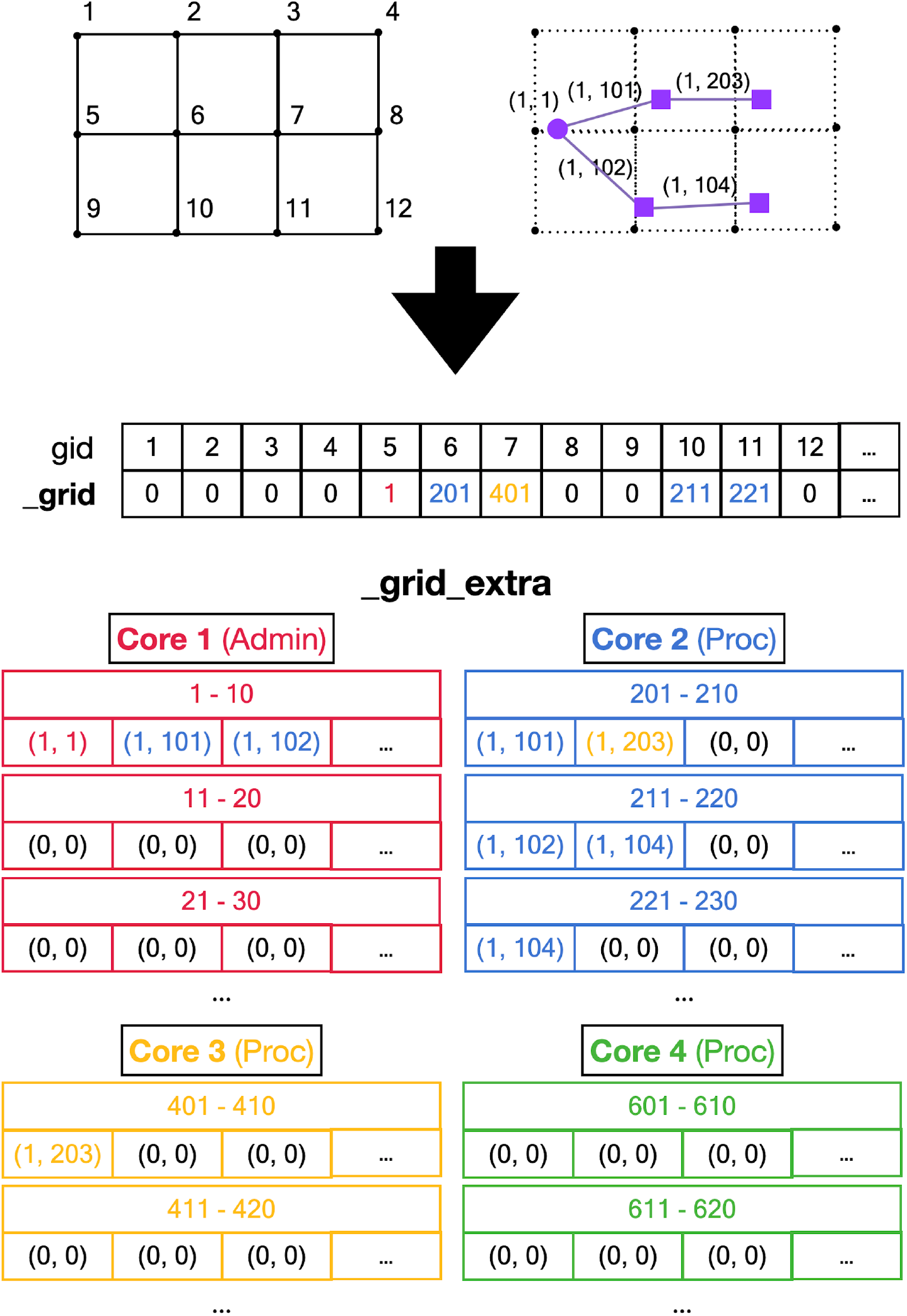
Registering front coordinates in a grid. Top: 2D representation of different grid locations in space, in actual code the grid is 3D. Left shows the index *gid* associated with each location, right shows the purple neuron from Figure 4 mapped onto the grid with front IDs. **Mid**: `_grid` shared array is indexed by *gid*. If no fronts are located at a grid point the entry is zero, otherwise it is an index into the `_grid_extra` array. Used indices are color coded for the Core that made the entry. **Bottom**: representation of `_grid_extra` shared array. This linear array is subdivided in private sections for each Core, color coded. When a Core needs to register a coordinate at a grid location that is empty it will allocated a new block (default size 10) in its private section of `_grid_extra`. For example, when `Admin_agent` (Core 1) makes soma (1,1) during the `add_neurons` call it allocates a block for _`grid` point 5 starting at index 1 in `_grid_extra` and writes the front ID (color coded for the Core that made the entry) at that index. As shown in Figure 4, Core 2 makes two cylindrical child fronts for (1,1) during cycle 1. Because the origin coordinates of these fronts (1,101) and (1,102) are located close to _`grid` point 5, they are entered at the next free indices 2 and 3 in the red block in `_grid_extra`. The end coordinates of (1,101) and (1,102) are close to _`grid` points 6 and 10, respectively. So Core 2 allocates two new blocks in its private section of `_grid_extra`, starting at indices 201 and 211, and enters the corresponding front IDs. During cycle 2 front (1,203) is created by Core 3, which allocates a new `_grid_extra` block at index 401 for _`grid` point 7, and front (104) is created by Core 2, which allocates a new `_grid_extra` block at index 221 for _`grid` point 11. The origin coordinates of these two fronts are added to previously allocated blocks at indices 202 and 212. When blocks fill up, for example when an 11th front needs to be allocated to _`grid` point 5 using indices 1-10 in `_grid_extra`, a new block needs to be allocated in `_grid_extra` (not shown), with for this example indices 421-430. This new block is linked to the existing one by changing the entry at index 10 to (−421,0) and the front ID that used to at index 10 is moved to index 421 and the the 11th front is registered at index 422.

This is achieved by a layered locking mechanism illustrated in Figures 6 and 7. Each core is identified by their number, called *pid*. They have a private location in `_grlock_request` and in `_gwlock_request` arrays in shared memory, which by default contain zeros. When they need access to the 3D grid they put the corresponding *gid* value in their private location of one of these request arrays and wait till the lock is obtained or a predetermined period has passed. If a lock cannot be achieved in time, a `GridCompetitionError` is raised. The request arrays are monitored by the `_lock_broker` method that is run intermittently by `Admin_Agent`. Because `_lock_broker` is the only method that is allowed to set grid locks, locking is effectively a serial process.

**Figure 6:**
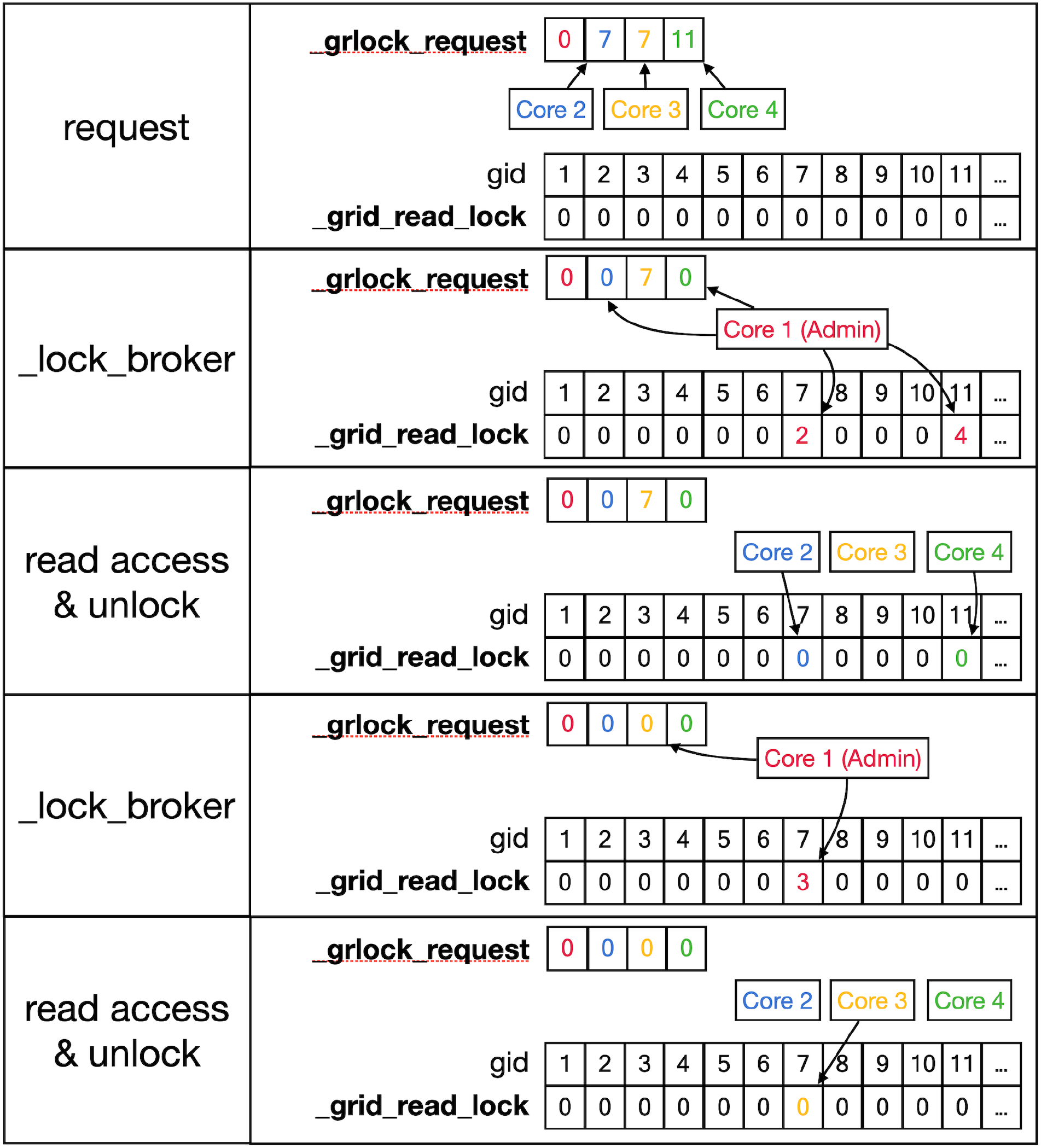
Sequence to lock _grid arrays for reading. Shows a series of events resulting in giving a Core read access to the `_grid` and `_grid_extra` shared arrays. **Top row**: As described in text, Cores write *gid*, an index in to the _`grid` array (see Figure 5) into their private location in the shared `_grlock_request` array, *gids* are color coded for Cores. Cores 2 and 3 both request access to *gid* 7, Core 4 requests access to *gid* 11. The `_grid_read_lock` shared array indicates the locking status for each *gid* value, with zero meaning no lock. Colored entries in this array will be coded for the Core that did the writing. **Second row**: `_lock_broker`, a method running serially on Core 1, reads entries in `_grlock_request` and locks the corresponding *gid* locations in `_grid_read_lock` if they were zero by entering the number of the core. It also resets the request in `_grlock_request` to zero. Because both Cores 2 and 3 requested access to *gid* 7, only Core 2 is giving a lock and Core 3 has to wait. **Third row**: Cores 2 and 4 notice that their indices appeared in `_grid_read_lock` for the *gid* entries they requested (7 and 11 respectively) and they now read information in the grid arrays, when done they clear the lock in `_grid_read_lock` by setting it back to zero. The request of Core 3 in `_grlock_request` remains active. **Fourth row**: `_lock_broker` can now execute the request of Core 3 and locks *gid* location 7 in `_grid_read_lock` for Core 3, again resetting `_grlock_request`. **Bottom row**: Core 3 can now perform a grid read at *gid* 7 and unlocks `_grlock_request`.

**Figure 7:**
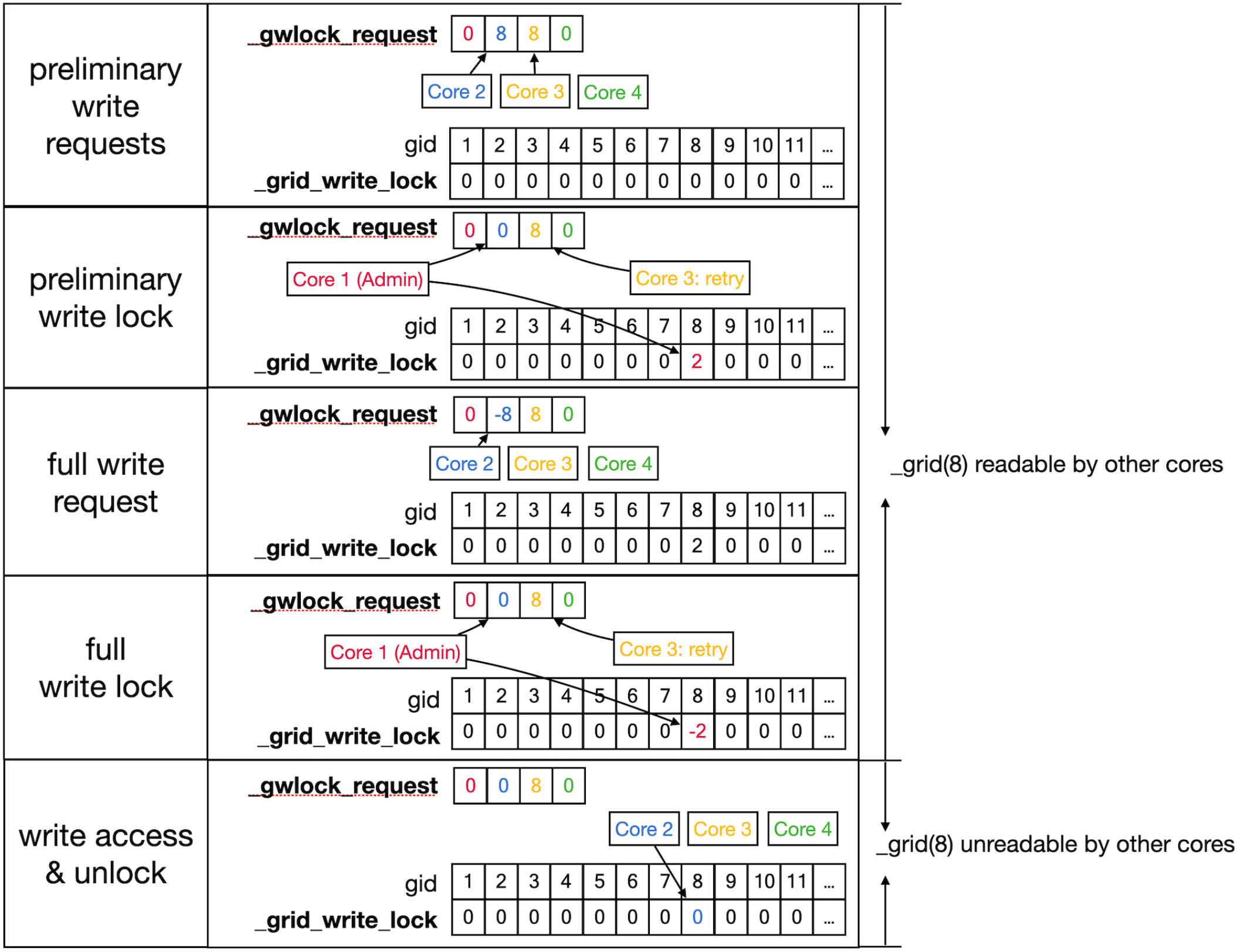
Sequence to lock grid arrays for writing. Writing access to the `_grid` and `_grid_extra` shared arrays uses shared arrays similar to those shown in Figure 6 for reading, called `_gwlock_request` and `_grid_write_lock`. Writing locks occur in two stages: the preliminary stage indicates that a Core may want to write to this location in the near future. The preliminary write locks are used for front scheduling by `Admin_agent` and during collision detection. During preliminary write lock of a *gid*, read locks can be obtained to it (see Figure 6). **Top row**: Core 2 and 3 both request preliminary write access to *gid* 8 in `_gwlock_request`. **Second row**: `_lock_broker` detects these requests and gives Core 2 preliminary write access by writing its index at *gid* 8 in `_grid_write_lock`. It also resets the request of Core 2, while that of Core 3 remains. **Third row**: sometime later Core 2 requests full write access by entering in `_gwlock_request` a negative *gid* value (−8) for the location it has already preliminary locked. Instead, if for some reason Core 2 notices that it does not requires writing, it will immediately remove the preliminary lock at *gid* 8 (not shown). The preliminary write request of Core 3 is still waiting. **Fourth row**: `_lock_broker` gives write lock access by entering the negative index of the requesting Core (−2) in `_grid_write_lock` and clears `_gwlock_request`. **Bottom row**: noticing the -2 value at gid location 8 in `_grid_write_lock`, Core 2 now performs its writing action upon the grid arrays. During this write action no other cores are allowed read or write access to the corresponding memory locations. When done, Core 2 clears the lock. If the entire sequence described above took less than a second, Core 3 will now receive preliminary write access. If not Core 3 will raise a `GridCompetitionError`.

As shown in Figures 6 and 7, `_lock_broker` uses shared memory `_grid_read_lock` and `_grid_write_lock` arrays to lock specific access to gid entries in `_grid`. When `_lock_broker` detects a request it checks whether that gid is currently locked for the read (Figure 6) or write (Figure 7) action requested. If the lock value was zero it was unlocked and will be locked by setting it to the pid of the requesting Core, `_lock_broker` also clears the request in `_grlock_request` or `_gwlock_request` by setting it back to zero. The requesting Core monitors the value of `_grid_read_lock` or `_grid_write_lock` at index gid and starts accessing the grid once it is set to its unique pid value. When it finishes accessing the grid it resets the lock to zero. Note that cores can only access their private sections of the `_grlock_request` and `_gwlock_request` arrays, but they can access any index in the `_grid_read_lock` and `_grid_write_lock` arrays. Conversely, `_lock_broker` has access to all data structures shown.

The locking sequence described applies to locking for read only access (Figure 6), for writing it is more complicated (Figure 7). A preliminary write lock is obtained first, with a positive gid value in `_gwlock_request` array, followed by a full write block, with a negative gid value in `_gwlock_request` array, before the actual writing starts. Read locks can be issued for *gid* that are under preliminary write block, but not for *gid* that are under full write block because the memory content is unreliable during writing. The preliminary write lock is used to prevent different processing cores of trying to perform actions that could lead to attempted updates of identical grid points. During an `add_child` sequence, the grid points to be occupied by the prospective new front are computed and preliminary write locks are requested. If a lock cannot be obtained within 1 second a `GridCompetitionError` is raised, this is in general due to multiple cores trying to grow in the same area of simulation space. To reduce the likelihood of this happening, `Admin_agent` sets preliminary write locks for all grid points associated with a scheduled front and only schedules fronts for processing that do not occupy already write locked grid points. The main customer of grid locking is the `_test_collision` method that checks whether proposed new fronts collide with existing ones. It is highly optimized to, if necessary, try obtaining needed preliminary write locks or read locks repeatedly to reduce the likelihood of `GridCompetitionError`. If the new front passes the collision check, it will be added to the grid and for that the full write lock has to be obtained first, but this lock is maintained only very briefly.

It is up to the user how to deal with a `GridCompetitionError`. Often the `add_child` method causing it is just repeated in the hope that other cores will release their lock on the grid points needed. But this should be tried only a few times to avoid wasting time on persistent `GridCompetitionErrors`. In case of simulations with many cycles, a good alternative is to abort the `add_child` call and try again during the next cycle.

## Results

The results section of this paper will focus on the performance of the software. Its practical use in simulating complex models of neural development is demonstrated in Kato and De Schutter (2022), a simulation of 10 postnatal days of the massive migration of cerebellar granule cells from the top of the simulation volume to the granular layer at the bottom, combined with the initial growth and partial retraction of Purkinje cell dendrites. The model contains more than 3000 neurons.

First performance on a performant laptop is described because many small models can be run effectively on such accessible hardware. Next extensive benchmarking is described, including comparisons with NeuroMaC. These benchmarks were run on a dedicated PC with a 32 core CPU.

### Running NeuroDevSim on a laptop

For simple simulations, like those in the notebooks in the examples directory, run time is quite short on a modern laptop. For example, approximate run times on 4 cores on a laptop with several other apps open are respectively 3.6 - 6.0 s, 2.3 - 3.2 s and 23 - 52 s for the models shown in Figures 1B-D with live plotting enabled (n = 5). Plotting strongly increases run times, without plotting run times become 0.5 - 0.9 s, 0.3 - 1.0 s and 1.3 - 1.9 s respectively.

Note that run times are quite variable. A systematic analysis of run times and its dependence on the number of cores used was done for the L5 pyramidal neuron model, simulating just one cell (Figure 8A). As expected mean run times rapidly decrease with increasing number of cores, speeding up from 9.8 s for serial simulation (2 cores) to 3.3 s for parallel simulation with 6 cores. Beyond 6 cores there is no further speed-up, instead run times increase slightly. This behavior is due to the small model size, as will be shown in a later section were a much larger model is benchmarked.

**Figure 8:**
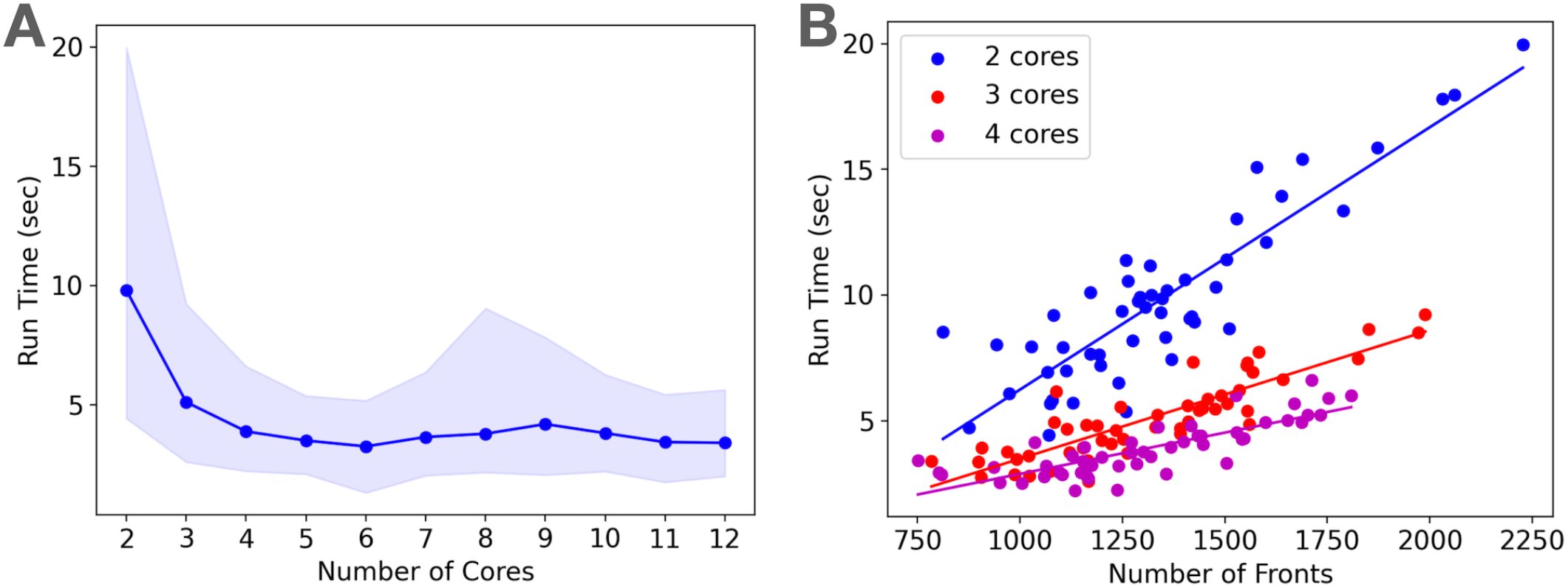
Run times of L5 pyramidal neuron model on a MacBook Pro laptop. (**A**) Mean run time in seconds for 120 cycles of simulation versus number of cores used (n = 50). Shaded area plots minimum and maximum run times, showing a large variance in run times that decreases with number of cores used. Note that the minimum number of cores is 2: one for *Admin_agent* and one for *Proc_agent*, resulting in a serial simulation. All other core numbers are for parallel simulation, with (number of cores - 1) processing cores. Run times rapidly decrease up to 5 processing cores (6 cores total), parallel simulation is less efficient for more cores due to the small model size. The hardware had 8 high performance cores. (**B**) Single trial run time in seconds versus number of fronts generated by the simulation for different number of cores used. Variability of run times was strongly correlated with the number of fronts generated. Correlation coefficients were 0.93, 0.93 and 0.82 for 2, 3 and 4 cores. For all cases in panel A the correlation was highly significant with p < 0.0001.

Run times were quite variable from simulation to simulation, as shown by the shaded area in Figure 8A. The variability of run time was strongly correlated with the number of fronts generated during the simulation (Figure 8B). Because branching events are probabilistic in this model, the number of branching events varies between simulation runs resulting in small models with few branch points or large ones with many branch points, depending on the random number sequence. As the number of cores increases, the highly variable number of fronts generated on each core is averaged by a larger n, resulting in lower variance of the total number of fronts and somewhat less variable run times.

Importantly, each parallel NeuroDevSim simulation is always unique, it cannot be reproduced for reasons described in Materials and Methods. In fact, the random seed which can be specified during `Admin_agent` instantiation only affects the random placement of somata by the `add_neurons` method. To obtain a reproducible simulation NeuroDevSim has to be run in serial mode with only a single processing core (2 cores in total), only then will the random seed also affect the total simulation outcome. The user has the choice between slow, reproducible serial simulation or fast, but unreproducible, parallel simulation. For model debugging purposes serial simulation may be preferred, but otherwise the speed advantages of parallel simulation are obvious.

### Memory Use

NeuroDevSim default settings allow for simulations with up to 20,000 fronts, which enables modeling the growth of hundreds of small neurons or a few large ones. Default memory use is 96 MB for simulations using a few cores, this increases to up to 221 MB for simulations using more than 10 cores. For larger simulations like the forest of 100 layer 5 pyramidal neurons simulation shown in Figure 1A that is used for benchmarking, two array sizes had to be increased to accommodate up to 30,000 fronts. This increased memory use to 209 MB for 2 core simulations, 1.1 GB for 10 core simulations and 1.7 GB for 25 to 32 core simulations.

### Performance benchmarks

NeuroDevSim run times for different model sizes were evaluated on a dedicated PC with a 32 core CPU and compared to similar benchmarks for NeuroMaC. For these benchmarks, run times of 10 simulations of 115 cycles each of the forest of layer 5 pyramidal neurons simulation shown in Figure 1A were averaged (Figure 9). The number of fronts made by NeuroMaC was also highly variable and scaled with run time. Minimum - maximum range over all simulations was slightly smaller for NeuroMaC (126,655 - 137,083 fronts) than for NeuroDevSim (124,013 - 138,997 fronts) but because less NeuroMaC simulations were run this small difference may be due to under sampling.

**Figure 9:**
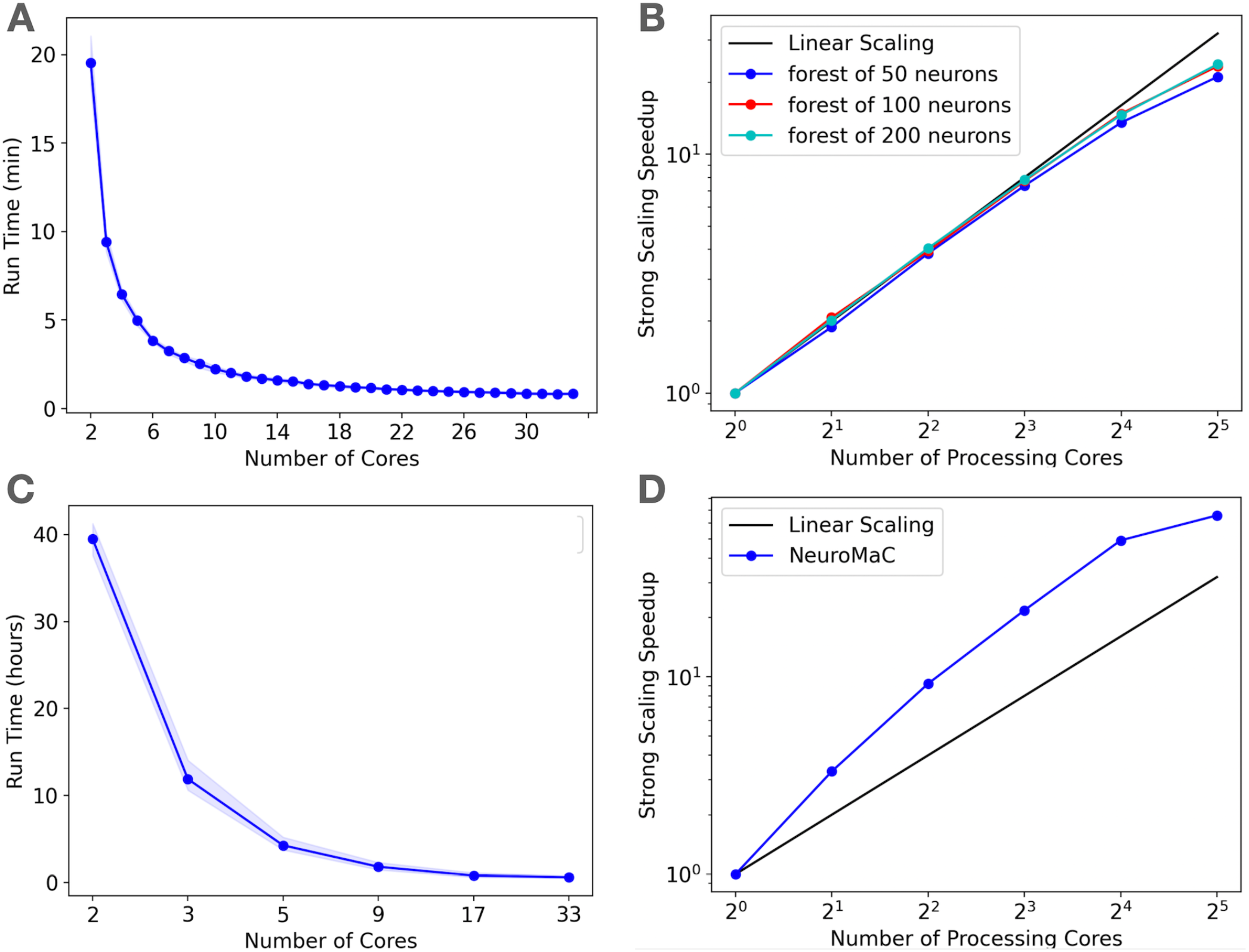
Run times of L5 pyramidal forest model using NeuroDevSim (A-B) or NeuroMaC (C-D) on AMD Ryzen Threadripper 32 core CPU. (**A**) Mean run time in minutes for 115 cycles of NeuroDevSim simulation of the L5 pyramidal forest of 100 neurons versus number of cores used (n = 10). (**B**) Strong scaling of NeuroDevSim for 3 different forest sizes of 50, 100 and 200 neurons respectively. Note that scaling improves with forest size, though differences between 100 and 200 neuron forests are small. (**C**) Mean run time in hours for 115 cycles of NeuroMaC simulation of the L5 pyramidal forest of 100 neurons versus number of cores used (n = 10). Note that the available number of processing cores combinations is limited, they correspond to spatial (x, y, z) subdivisions of (1,1,1), (2,1,1), (4,1,1), (8,1,1), (8,2,1), (16,2,1) respectively. Other subdivision schemes, like (4,2,1) and (4,4,1), were also tried but gave similar results. Shaded area plots minimum and maximum run times. (**D**) Strong scaling of NeuroMaC for the simulations shown in **C**.

For the standard simulation of a forest of 100 neurons, NeuroDevSim run times varied between 19.5 minutes for serial simulation and 0.84 minutes for 32 core simulation (Figure 9A). Start up times were negligible at 0.08 seconds for serial simulation, increasing to 0.10 seconds for 32 core simulation. This compares to much slower NeuroMaC run times for the same model, varying between 39.5 hours for serial simulation and 36.0 minutes for 32 core simulation (Figure 9C) with a close to constant start up time of about 2 seconds. In conclusion, NeuroDevSim simulations are about 2 orders of magnitude faster than NeuroMaC simulations, confirming the speed ups obtained from using shared memory parallel simulation with continuous scheduling of front processing compared to a messaging based spatial parallelization.

How effective is the parallelization? This was evaluated by computing strong scaling, where linear performance corresponds to a reduction of run time by a factor two for doubling the number of processing cores. As shown in Figure 9B, NeuroDevSim achieves close to optimal scaling for up to 8 cores, but for 16 and 32 cores there is a reduced scalability. This depends on the size of the model simulated, with a smaller loss of scalability for the 200 neuron forest than for the 50 or 100 neuron ones. This strongly suggests that the scaling problem is caused by the end of each simulation cycle, when an increasing number of cores become idle while the last few fronts are processed by a subset of cores. The total simulation run times scale close to linear with forest size: for the 50 neuron forest 9.2 minutes for serial simulation and 0.43 minutes for 32 cores, and respectively 39.2 minutes and 1.65 minutes for the 200 neuron forest. The more effective strong scaling for the larger forest is therefore due to the idle time at the end of each simulation cycle becoming proportionally shorter. Using hyperthreading to simulate with 64 processing cores was not effective (not shown), this was expected because the physical cores are kept quite busy.

Another factor affecting scaling for large number of cores might have been delays during `_lock_broker` calls (Figures 6, 7). Because `_lock_broker` is run intermittently on the `Admin_agent` core, there may be short wait times before it responds and this might scale with the number of processing cores. Therefore, a separate series of benchmarks was run where `_lock_broker` ran continuously on a separate core. This did not result in any speed-up (not shown) and because it used an extra core was not an effective parallelization strategy.

Finally, strong scaling was also evaluated for NeuroMaC (Figure 7D). Here performance was initially supra linear, with a big speed increase in going from serial to parallel with just 2 processing cores. But this effect only lasted up to 16 processing cores, with comparatively little gain for 32 cores.

## Discussion

This paper describes NeuroDevSim, to our knowledge at present the only actively maintained simulator of neural development that can model migration of neurons, growth of dendrites and axons and formation of early synaptic connections. In this regard, NeuroDevSim replicates most of the functionality of the no longer maintained CX3D (Zubler, Hauri et al. 2013) with the exception of progenitor cell proliferation. Though birth of new neurons can be made conditional in NeuroDevSim by adding the relevant statements before the `add_neurons` method (Figure 2), modeling actual neural proliferation is a goal for future NeuroDevSim versions. In practice, simulation of neural proliferation is rather artificial in a fixed simulation volume, as used in both CX3D and NeuroDevSim. Brain regions undergoing large proliferation rapidly expand in volume (e.g. (Legue, Riedel et al. 2015) and not simulating this expansion leads to artificially increased crowding conditions in the model. In our simulation of cerebellar development this was identified as the major limitation of the model (Kato and De Schutter 2022). Nevertheless, as shown in Zubler, Hair et al (2013), in Torben-Nielsen and De Schutter (2014) and in Kato and De Schutter (2022) highly detailed models of neural tissue development can be simulated with these simulators.

Besides enabling simulation of neural development, NeuroDevSim also introduces an innovative way of parallelizing computation based solely on using shared memory in multi-core architectures. The number of cores on common CPUs has increased rapidly and our approach can now be used with up to 128 cores on platforms that combine two 64 core AMD CPUs. We have successfully simulated the Kato and De Schutter (2022) model on 128 cores, though as the speed-up was sublinear we decide to run it on 64 cores for that study. Because there are no communication delays in shared memory simulation, it outperforms messaging based parallelization spectacularly as demonstrated in the comparison of NeuroDevSim to NeuroMaC run times (minutes versus hours in Figure 9) and achieves close to optimal parallelization for up to 16 cores for the models used in this study. Based on our experience in Kato and De Schutter (2022) better scaling performance for 32 cores is expected for larger and more complex models than the forests simulated here. As it is likely that future CPUs will offer increasing numbers of cores, NeuroDevSim can be expected to simulate even larger and more complex models.

Implementation of shared memory parallelization introduces unique programming challenges: the use of fixed size data arrays and the need to avoid memory write conflicts. Though not explored in this study, we expect that these challenges are fairly easy to solve for fixed data size problems. The size of data arrays is predictable for such applications and the use of private sections of data arrays for writing as demonstrated in Figures 3 and 4 should be sufficient to avoid memory conflicts. Unfortunately, this did not apply to NeuroDevSim because array sizes are not predictable when modeling stochastic growth. As a consequence array sizes need to be estimated and, in practice, this is often based on trial and error. If NeuroDevSim models become too large for the default array size parameters, the simulation will crash with an `OverflowError` that mentions which array size parameter needs to be increased. In the absence of `OverflowError`, the allocated arrays are often too large but with CPU memory use in the range of 200 MB to 2 GB this should not be a concern on most computing platforms.

With this caveat, most of data allocation and updating could be handled in a rigorous way that avoids memory conflicts, except for collision detection. Collision avoidance is a tough challenge in parallel computing, irrespective of whether messaging or shared memory approaches are used. Solving the conflict of new structures being generated on different cores that overlap in physical space requires, by definition, interaction between the computing cores involved. In the case of messaging based parallelization this requires first communication to detect a possible collision and in case of collision extra messaging to solve it, the latter may be challenging if more than two cores are involved. In the case of our implementation of shared memory parallelization (Figure 5), detection is fast but solving the collision is less tractable because cores do not communicate directly. Therefore the simulator tries to avoid these problems as much as possible by not scheduling simultaneous updating of fronts that occupy nearby areas of simulation space. As the number of cores used increases this may, however, become infeasible leading to `GridCompetitionError` on one of the cores involved and requiring the user to code solutions for such occurrences.

In fact, besides programming challenges, shared memory computation requires the software user to follow a set of strict rules that our outlined in the NeuroDevSim documentation. Unusual for Python programs, instance attributes cannot be used and many front or synapse attributes should never be accessed or changed directly, instead specific methods have to be called. Similarly, fronts or synapses can only be made using NeuroDevSim methods, they cannot be instantiated by the user. And, as already mentioned, the user may have to change preset array sizes during the `Admin_agent` instantiation and have to adapt the model code to deal with `GridCompetitionError`. To help users in learning how to deal with these challenges many coding examples are provided and, once one gets used to the specific coding style, NeuroDevSim models can be written in a clear and concise style.

NeuroDevSim 1.0 is a first version of the software. As we gain more experience in using it to simulate real development, additional features and improvements will be added. An obvious priority is to deal with the fixed simulation volume issue. However, extensive research is needed to decide which of several possible approaches will result in the biologically most realistic outcome.

## Acknowledgements

This work was performed with funding from the Okinawa Institute of Science and Technology Graduate University. The author wishes to thank Dr. Weiliang Chen for help with the Figures and Drs. Weiliang Chen, Iain Hepburn and Jules Lallouette for helpful comments on previous versions of this manuscript. Mizuki Kato acted as tester of preliminary versions of the software.

## Notes

https://github.com/CNS-OIST/NeuroDevSim

### Competing Interest Statement

The authors have declared no competing interest.

## Bibliography

Ascoli, G. A., J. L. Krichmar, S. J. Nasuto and S. L. Senft (2001). “Generation, description and storage of dendritic morphology data.” Philosophical transactions of the Royal Society of London Series B, Biological sciences 356(1412): 1131–1145.

Bower, J. M. and D. Beeman (1998). The book of GENESIS: exploring realistic neural models with the GEneral NEural SImulation System, pringer-Verlag New York.

Goodman, D. F. and R. Brette (2009). “The Brian simulator.” Front Neurosci 3(2): 192–197.

Hines, M. L. and T. T. Carnevale (1997). “The NEURON simulation environment.” Neural computation 9(6): 1179–1209.

Jordan, J., T. Ippen, M. Helias, I. Kitayama, M. Sato, J. Igarashi, M. Diesmann and S. Kunkel (2018). “Extremely Scalable Spiking Neuronal Network Simulation Code: From Laptops to Exascale Computers.” Front Neuroinform 12: 2.

Kato, M. and E. De Schutter (2022). “Granule cells affect dendritic tree selection in Purkinje cells during cerebellar development.” In preparation.

Koene, R. A., B. Tijms, P. van Hees, F. Postma, A. de Ridder, G. J. A. Ramakers, J. van Pelt and A. van Ooyen (2009). “NETMORPH: a framework for the stochastic generation of large scale neuronal networks with realistic neuron morphologies.” Neuroinformatics 7(3): 195–210.

Legue, E., E. Riedel and A. L. Joyner (2015). “Clonal analysis reveals granule cell behaviors and compartmentalization that determine the folded morphology of the cerebellum.” Development 142(9): 1661–1671.

Sanes, D. H., T. A. Reh, W. A. Harris and M. Landgraf (2019). Development of the Nervous System, Academic Press.

Torben-Nielsen, B. and E. De Schutter (2014). “Context-aware modeling of neuronal morphologies.” Frontiers in neuroanatomy 8: 92.

van Ooyen, A., Ed. (2003). Modeling Neural Development, MIT Press.

Zubler, F. and R. J. Douglas (2009). “A framework for modeling the growth and development of neurons and networks.” Frontiers in Computational Neuroscience 3: 25.

Zubler, F., A. Hauri, S. Pfister, R. Bauer, J. C. Anderson, A. M. Whatley and R. J. Douglas (2013). “Simulating Cortical Development as a Self Constructing Process: A Novel Multi-Scale Approach Combining Molecular and Physical Aspects.” PLoS Computational Biology 9(8): e1003173.

